# Subindividual variation in plants: branch-level phenotypic and epigenetic divergence of pollen grains is correlated with the epigenetic divergence of adjacent leaves

**DOI:** 10.64898/2026.06.13.732073

**Authors:** Carlos M. Herrera, Mónica Medrano, Pilar Bazaga, Conchita Alonso

## Abstract

The persistence across generations of somatic epigenetic mosaicism arising within plants from random drift or localized responses to the environment is an important determinant of the potential evolutionary significance of subindividual plant variation, yet practical limitations have so far precluded its study in natural plant populations. We exemplify here an indirect approach based on testing whether branch-level somatic epigenetic mosaicism exhibited by individuals of a long-lived tree (diploid sporophytes) is consistent with the phenotypic and epigenetic mosaicism exhibited by the pollen grains (haploid gametophytes) produced by the same branches.
Paired samples of leaves and pollen from five distinct branches within each of five wild-growing *Pistacia terebinthus* L. (Anacardiaceae) trees were characterized epigenetically using genome-wide DNA cytosine methylation level and multivariate epigenetic fingerprinting. Pollen was also characterized phenotypically by grain dimensions and shape.
Individual trees were somatically heterogeneous with regard to cytosine methylation level and multivariate epigenetic fingerprints of leaf tissue. Phenotypic and epigenetic features of pollen grains differed significantly among branches of the same tree. Intraplant variation in pollen phenotype and multivariate epigenetic fingerprints were significantly related to intraplant variation in epigenetic leaf features.
The consistency of subindividual somatic epigenetic heterogeneity of leaves with epigenetic and phenotypic heterogeneity of haploid gametophytes from the same branches suggests that epigenetic variants arising in somatic tissues over individual lifespans could eventually contribute to the phenotypic diversity represented in the next adult generation and become established in populations. This mechanism might contribute to enhance epigenetic and phenotypic diversity in natural plant populations.

**One-sentence summary:** Pollen grains produced by differente branches of the same tree are heterogeneous with respect to size measurements and epigenetic profiles, and such subindividual heterogeneity is correlated with subindividual variation in the epigenetic profiles of the leaves in the same branches.

## Introduction

The modular construction of plants, involving the proliferation of nonidentical homologous structures, gives rise to a distinct subindividual component of phenotypic and epigenetic variance in natural plant populations (Alonso et al., 2018; Herrera et al., 2022; Yao et al., 2021). In contrast with a tradition that negates ecological or evolutionary relevance to the variation that occurs within genetically homogeneous individuals (Mayr, 1982; Herrera, 2024), recent observational and experimental research has shown that such variation can actually possess some ecological significance. For example, subindividual phenotypic variability is responsive to the environment (Pélabon et al., 2021; Møller et al., 2023; Castro Sánchez-Bermejo et al., 2025), broadens the functional niche of individuals and populations (Alonso et al., 2018; Herrera et al., 2015, 2019, 2021, 2022), influences the outcome of interactions with animals (Benitez-Vieyra et al., 2014; Shimada et al., 2015; Wetzel et al., 2016), and modulates the effects of phenology, mating system and floral traits on individual fitness (Arceo-Gómez et al., 2017; Maldonado et al., 2023; Paglia et al., 2023; Ehrlén & Valdés, 2024; Trevizan et al. 2024, 2026). The possible evolutionary significance of subindividual variability, however, is still largely unknown. Epigenetic marks which often underly subindividual phenotypic variation (Das & Messing, 1994; Bitonti et al., 1996; Gao et al., 2010; Herrera & Bazaga, 2013; Alonso et al., 2018; Herrera et al., 2022) may or may not persist unchanged after gametogenesis and zygote formation (Richards, 2006; Akimoto et al., 2007; Jablonka & Raz, 2009; Bošković & Rando, 2018). Evolutionary consequences will therefore depend on the inheritance of intraplant epigenetic variants, for roughly the same reasons ordinarily argued in relation to the evolutionary significance of somatic mutations that involve modifications in DNA sequence (Schoen & Schultz, 2019; Lesaffre, 2021; Schmitt et al., 2024). Conceptual and practical difficulties, however, have so far hindered progress along that line of inquiry. On one hand, the application of quantitative genetics models to assess inheritance of “softly heritable” epigenetic features is conceptually problematic (Gorelick, 2005; Johannes et al., 2008; Banta & Richards, 2018). And on the other hand, formal inheritance studies are impractical with long-lived plants, which are precisely those where epigenetic mosaicism should be most frequent because of a combination of high spontaneous epimutation rates, huge number of lifetime cell divisions and extensive propagation of epimutated cell lineages during the ontogenetic unfolding of shoot branching (Lesaffre, 2021; Herrera et al., 2021; Chen et al., 2024; Johannes, 2025; van der Graaf et al., 2025).

Examining whether epigenetically divergent parts of a given adult plant (diploid sporophyte) produce phenotypically and/or epigenetically divergent populations of haploid gametophytes (pollen or ovules) provides an indirect pathway to explore the potential evolutionary significance of epigenetically-based subindividual variation. This approach is motivated by the key precondition that, to be evolutionarily consequential, epigenetic divergence among different parts of the same sporophyte, either triggered by the environment or arising from somatic epigenetic drift (Herrera et al., 2021; Yao et al., 2021, 2023; Chen et al., 2024), should not be entirely wiped out by the epigenetic reprogramming that takes place during gametogenesis (Migicovsky & Kovalchuk, 2012; Gutierrez-Marcos & Dickinson, 2012). By adopting the preceding procedure, this paper examines whether somatic epigenetic variation occurring within reproductive individuals of the long-lived tree *Pistacia terebinthus*L. (Anacardiaceae) is consistent with subindividual variation in epigenetic and phenotypic features of male gametophytes. The following specific questions will be addressed: (1) Do distinct branches of individual trees differ in somatic epigenetic features ?; (2) Is the pollen produced by epigenetically distinct branches of the same tree significantly heterogeneous in phenotypic and epigenetic traits?; and (3) Is within-plant divergence in phenotypic and epigenetic traits of pollen concordant with somatic divergence among branches in epigenetic featuresQuestions (2) and (3) actually represent workable versions of the one which ultimately motivated this study, namely (4) Is epigenetic reprogramming during gametogenesis thorough enough as to produce an epigenetically homogeneous gametophyte population from a single plant? Pollen grain size and shape will be used here to phenotypically characterize the male gametophytes produced by different branches in the same tree, as intraspecific variation in these easily measured morphological traits can influence their performance (Cruzan, 1990; Kelly et al. 2002; McCallum & Chang, 2016). Estimation of genome-wide DNA cytosine methylation and multivariate epigenetic fingerprinting with anonymous methylation-sensitive markers (Herrera et al., 2021, 2022) will be used to epigenetically characterize branches of individual trees with regard to both somatic tissues (expanding new leaves) and gametophytes (pollen grains) being produced simultaneously.

## Materials and methods

### Study plant and field sampling methods

*Pistacia terebinthus* is a long-lived, deciduous, dioecious, wind-pollinated tree widely distributed in the Mediterranean Basin. In male individuals, clusters of new expanding leaves develop in spring at branch tips at the same time that adjacent inflorescences are shedding pollen (Figure 1). Simultaneous timing and spatial closeness of leaves and inflorescences make possible paired sampling of sporophytic and gametophytic tissues of identical age borne by the same growth axis and sharing the same developmental and genealogical history. Repeating the paired sampling scheme on different branches of the same crown thus allowed a concurrent assessment of within-plant epigenetic variability in fresh leaves and pollen while controlling for possible seasonal and age-related differences. Another advantage of *P. terebinthus* for addressing the questions considered in this study is that male inflorescences produce copious pollen, which made possible obtaining sufficient DNA for epigenetic fingerprinting and chemical analysis of cytosine methylation (see below).

**Figure 1.**
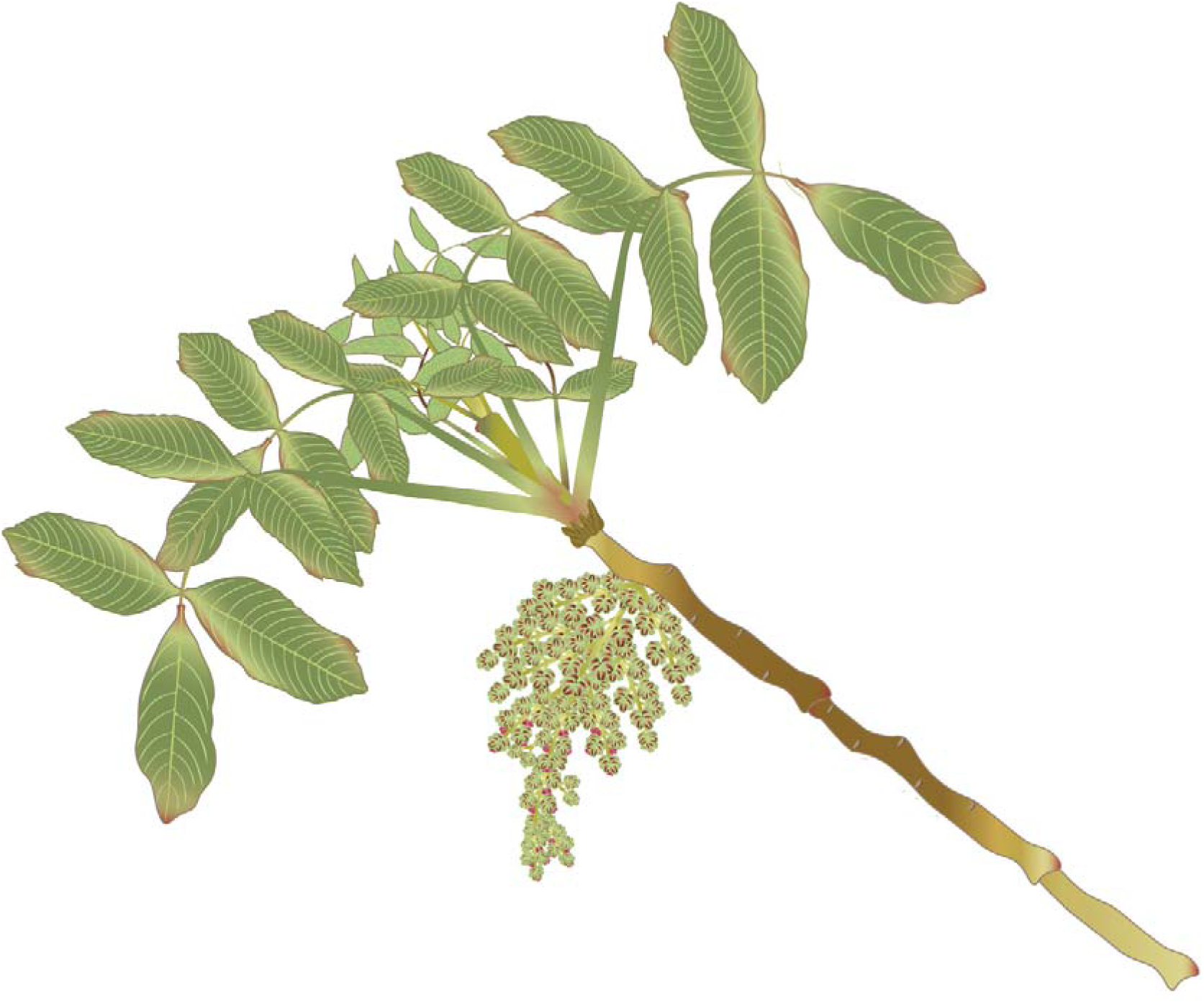
Representation of the tip of a branch of a male *Pistacia terebinthus* tree at leaf flushing time, to illustrate the close spatial proximity and temporal synchrony of expanding fresh leaves and pollen-shedding inflorescences, on which the paired leaf-pollen sampling scheme used in this study was based.

Five male *P. terebinthus* trees were chosen in a dolomitic outcrop at 1100 m a.s.l. in the Sierra de Cazorla, Jaén province, Spain. Within each tree, five paired samples each consisting of expanding fresh leaves and adjacent pollen-shedding inflorescences (Figure 1) were collected from the tip of major branches widely spaced in the crown. Leaf samples were placed inside paper bags, dried at ambient temperature in containers filled with silica gel, and kept at ambient temperature until processed. To promote anther dehiscence and pollen shedding, inflorescence samples were left to dry on aluminium foil sheets within containers with silica gel. Once anthers had released most pollen the aluminium foil was folded over itself, stored in a paper bag and kept dry at ambient temperature until processed.

## Laboratory methods

Dry leaf samples were homogenized to a fine powder using a Retsch MM 200 mill. Purified pollen samples were obtained by separating pollen grains from inflorescence remains (anther walls, inflorescence axes, bracts) following with minor modifications the procedure of Johnson-Brousseau and McCormick (2004), based on using different-pore sized Nitex® meshes (80, 38 and 6 µm pore size meshes in our case; Sefar America, Inc., Depew, NY, USA), held together in sequence and attached to a handheld vacuum cleaner. The 80 µm mesh filtered out most of the larger flower parts and debris of unwanted maternal tissue, while the 38 µm mesh trapped smaller debris and small amounts of pollen. Most of the pollen passed through these two first mesh filters and was trapped on the third, 6 µm mesh. Samples of dry material stored into the aluminium foil bags were released inside hand-made 80 µm mesh bag and immediately placed inside other two nested mesh bags (38 µm and 6 µm for inner and outer bags, respectively). To harvest the purified pollen sample, the set of three nested bags with decreasing pore size holding the plant material was placed inside a plastic plumbing tube attached to a handheld vacuum cleaner and exposed to the suction force of the device. A paintbrush was used to release the pollen gathered into the 6 µm filter, which was stored dry into a 1.5 mL microcentrifuge tube. This procedure was repeated on aliquots of the same field sample until sufficient quantity of pollen was obtained. On average, 125 mg of purified pollen was obtained from each field sample (range = 40-330 mg).

Total genomic DNA was extracted from leaf (*N* = 25) and pollen (*N* = 25) samples using the Omega Biotek E.Z.N.A. Plant DNA DS Mini Kit, and digested with DNA Degradase Plus (Zymo Research), a nuclease mix degrading DNA to its individual nucleoside components. Digested samples were stored at -20°C until analysed. To allow for properly testing the statistical significance of within-tree heterogeneity (i.e., among branches) in DNA methylation level of leaves and pollen, two (leaves) or six (pollen) aliquots were prepared from each DNA hydrolyzate, and the resulting *N* = 200 samples (5 plants x 5 branches x 8 aliquots) were processed in randomized order. Genome-wide percent methylation of DNA cytosines was determined for each sample with the chromatographic technique described by Alonso et al. (2016), using reversed phase HPLC with spectrofluorimetric detection. Global cytosine methylation was estimated for each sample as 100 x 5mdC/(5mdC + dC), where 5mdC and dC are the integrated areas under the peaks for 5-methyl-2’-deoxycytidine and 2’-deoxycytidine, respectively. The position of each nucleoside was determined using commercially available standards. Samples producing poor-quality DNA, weak chromatographic signal or unreliable peak area integration, were discarded. Results presented below are based on 45 (leaves) and 143 (pollen) independent HPLC runs.

Epigenetic fingerprinting of each leaf and pollen sample was conducted using a variant of the amplified fragment-length polymorphisms technique (AFLP) allowing to identify instances of intraplant polymorphism in methylation state of methylation-susceptible anonymous 5’-CCGG sequences. Since we were interested in assessing intraplant epigenotype heterogeneity rather than individual differences, our AFLP method used exclusively primer combinations based on the methylation-sensitive HpaII enzyme. HpaII cleaves 5’-CCGG sequences but is inactive when either or both cytosines are fully methylated. In absence of DNA sequence variation among samples, as expected for tissues from the same plant (estimates of plant somatic mutation rates ∼10^5^ times lower than somatic epimutation rates; Johannes 2025), any observed intraplant polymorphism in these methylation-sensitive AFLP markers (MS-AFLP hereafter) will reflect subindividual heterogeneity in the methylation state of the associated 5’-CCGG site (see Herrera et al., 2021, 2022 for previous applications of this simplified method). Leaf and pollen samples were fingerprinted with eight different primer combinations, each with two (HpaII) or three (MseI) selective nucleotides. Replicated analyses on different DNA aliquots (2-3 replicates per sample) were run for each leaf and pollen sample, leading to a total of 128 independent MS-AFLP runs for the 50 leaf and pollen DNA samples. A total of 450 MS-AFLP markers with variable methylation state were retained for analysis.

Variation in pollen grain size and shape was assessed by drawing aliquots from the same plant x branch pollen samples used for DNA extraction. Each pollen aliquot was evenly spread onto a microscope slide. Two drops of an aqueous solution of 0.01 % toluidine blue and a drop of 0.5 % glycerol were added to each slide and gently mixed with pollen grains. Grains thus stained were viewed using a Nikon Eclipse 80i microscope (Nikon instruments Inc. Melville, NY), and digital images of 10-15 randomly selected microscopic fields were taken for each individual slide using Nikon DS-U2-Ri1 camera and NIS-Elements software (version 3.22.15). Images were analyzed using ImageJ software (version 1.54p, Schneider et al., 2012). Each image was calibrated, converted to an 8-bit format, binarized using the auto threshold function with default parameters, and inverted to create a black background with pollen grains as white objects. The software functions erode, dilate, fill holes and watershed were used to delineate each object, fill blank spaces in the objects, and separate closely spaced ones. The “analyze particle” function was then used to measure the major and minor axes of the best-fitting ellipse for each particle, as well as to estimate its “circularity” (4·π·area/perimeter^2^, equalling 1.0 in the case of a perfect circle). A total of 13,110 pollen grains were measured, averaging 524 grains per plant x branch combination (interquartile range = 483-605 grains).

### Statistical analyses

Statistical analyses reported in this paper were carried out using the R environment (version 4.5.0; R Core Team, 2025). The same analytical procedure, based on fitting intercept-only, linear mixed-effect models to data using the lmer or glmer functions from package lme4 (Bates et al., 2015), was consistently adopted to estimate within- and among-tree variance in DNA methylation of leaf and pollen tissue, methylation status of polymorphic MS-AFLP markers, and pollen grains features. Tree and branch nested within tree were treated as random effects in all models to hierarchically partition total sample variance of response variables into among- and within-tree components. The null hypothesis that among-branch, within-plant variance of the response variable was not distinguishable from zero was tested by computing 95% confidence intervals for variance components using the confint.merMod function in lme4 with method = “boostrap” and 10,000 repetitions.

The hypothesis that within-tree somatic variation in epigenetic features was related to within-tree variation in phenotypic and epigenetic traits of the pollen was tested by fitting linear mixed-effects models with epigenetic leaf traits as predictors, and epigenetic and morphological pollen traits as response variables. Tree identity was included in all cases as a random effect, so that the predictive value of among-branch variation was tested in the within-plant context. Multivariate epigenetic fingerprints of leaves (predictors) and pollen (responses) were obtained as sample coordinates on axes from nonmetric multidimensional scaling (function metaMDS in package vegan; Oksanen et al., 2026) run on pairwise dissimilarity matrices computed from the binary matrix of methylation states for the *N* = 430 polymorphic MS-AFLP markers. To avoid artifacts and ensure *a priori* independence of the sets of coordinates for leaf and pollen samples, the conservative decision was adopted of running separate metaMDS for leaf and pollen data. The stress values obtained in these independent runs were 0.087 and 0.074 for leaves and pollen, respectively, denoting good ordinations of the multivariate epigenetic data and little risk of drawing false inferences (Dexter et al. 2028).

## RESULTS

### Somatic epigenetic divergence within trees

The two analytical methods used to characterize epigenetically the tissues of *P. terebinthus* trees yielded similar results when applied to fresh leaves, revealing that internal somatic epigenetic divergence within trees was greater than divergence among trees (Table 1A). Within-tree, among-branch variance in percent DNA cytosine methylation exceeded among-tree variance, and the lower limit of its 95% confidence interval was greater than zero (Table 1A). In contrast, the lower limit of the confidence interval for among-tree variance in cytosine methylation was zero, thus casting doubts on the significance of individual differences in DNA methylation level. When leaf tissue was characterized using multivariate analysis of the methylation status of anonymous MS-AFLP markers, estimated within-tree variance in per-marker methylation state likewise exceeded among-tree variance, and its confidence interval did not include zero. The confidence interval for the among-tree variance did include zero (Table 1A), again pointing to comparatively minor importance of somatic epigenetic variation among trees.

**Table 1.**
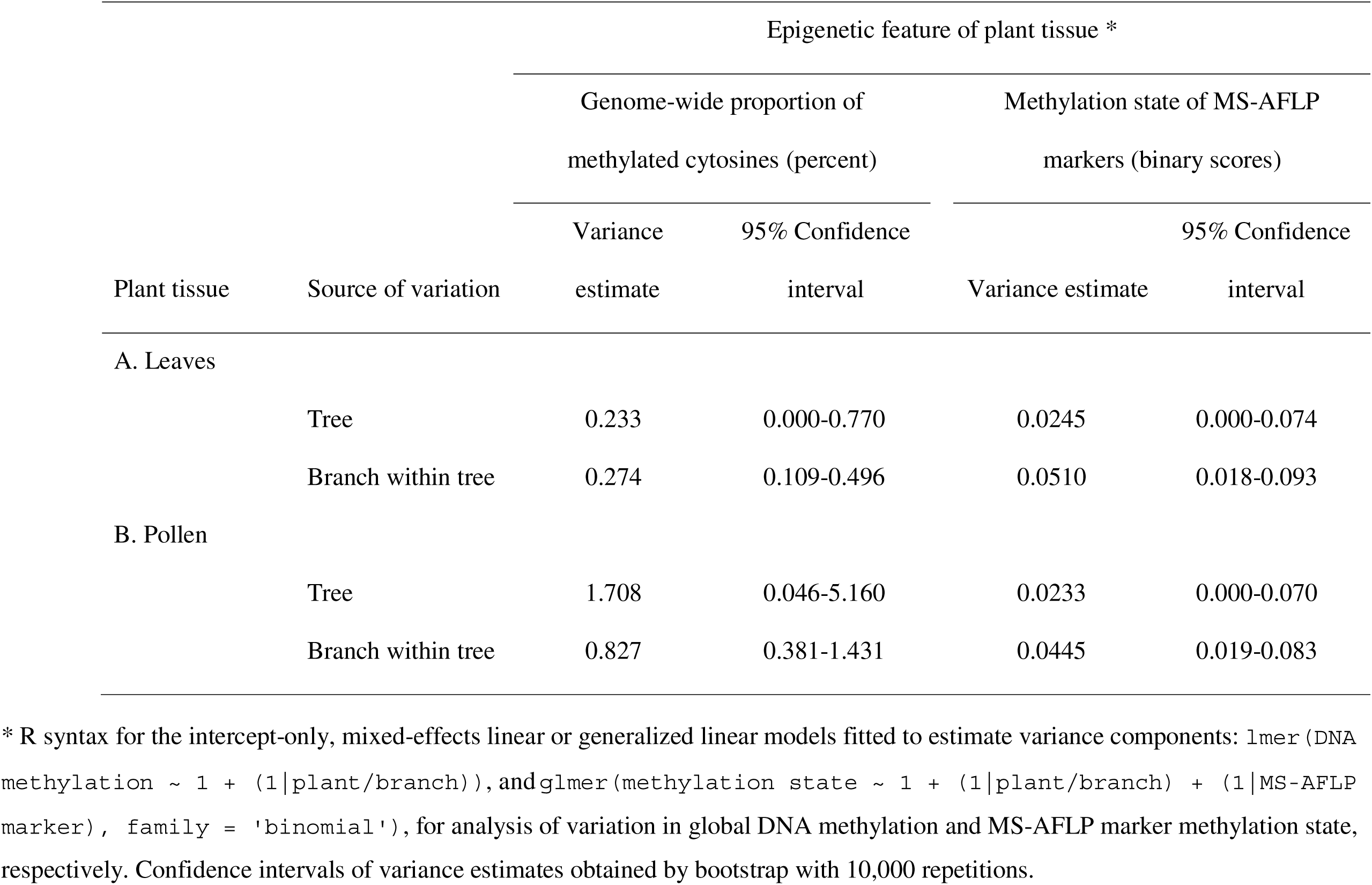
Among and within tree epigenetic variance components for leaf (A) and pollen (B) tissue.

### Pollen epigenetic and morphological divergence within trees

Pollen from different branches in the same tree differed widely in epigenetic features, as revealed by the two analytical procedures applied (Table 1B). DNA methylation and epigenetic fingerprints of the pollen produced by a given tree depended significantly on the branch where it was produced. The lower limit of confidence intervals for within-tree variance components were greater than zero for proportion of methylated cytosines and per-marker methylation state of MS-AFLP markers (Table 1B). The lower limit of the confidence intervals for among-tree variance estimates was zero for methylation state of MS-AFLP markers but not for the proportion of methylated cytosines (Table 1B).

Estimation of among- and within-tree variance components revealed substantial variation among branches of the same tree in pollen grain size (major and minor axis length) and shape (circularity) (Table 2). Among-tree variance estimates for the two linear dimensions were greater than within-tree variances, although these latter were still substantial and the lower limits of their confidence intervals were farther from zero than those for the confidence limits of among-tree variance estimates (Table 2). In the case of minor axis length and circularity measurements, the estimated confidence intervals for among-tree variance included zero (Table 2), further pointing to negligible individual variation in pollen grain size and shape in comparison to within-individual variation.

**Table 2.** Estimates of among and within tree components of variance for phenotypic traits of pollen grains.

| Source of variation | Phenotypic traits of pollen grains * |  |  |  |  |  |
| --- | --- | --- | --- | --- | --- | --- |
| | Major axis (µm) | | Minor axis (µm) | | Circularity ($4 \cdot \pi \cdot \text{area} / \text{perimeter}^2$) | |
|  | Variance | 95% Confidence | Variance | 95% Confidence | Variance | 95% Confidence |
|  | estimate | interval | estimate | interval | estimate | interval |
| Tree | 1.148 | 0.063-3.342 | 0.766 | 0.000-2.339 | 0.00094 | 0.000-0.0035 |
| Branch within tree | 0.412 | 0.196-0.709 | 0.512 | 0.239-0.883 | 0.00269 | 0.0013-0.0045 |

\* R syntax for the intercept-only, mixed-effects models fitted to estimate variance components: `lmer(pollen grain trait ~ 1 +` `(1 | plant/branch)` . Confidence intervals of variance estimates obtained by bootstrap with 10,000 repetitions

### Within-tree relationship between leaf and pollen epigenetic divergence

Differences among branches of the same tree in size and shape of pollen grains were significantly related to within-tree variation in multivariate epigenetic fingerprints of leaf tissue, but not to variation in genome-wide methylation level (Table 3). Across branches of the same tree, mean major and minor length of pollen grains were inversely related to scores on the third reduced axis of multivariate epigenetic variation of leaves () obtained from multidimensional scaling of the binary MS-AFLP matrix (Figure 2). The relationships between pollen dimensions and L-MDS3 were remarkably similar across the five trees sampled.

**Figure 2.**
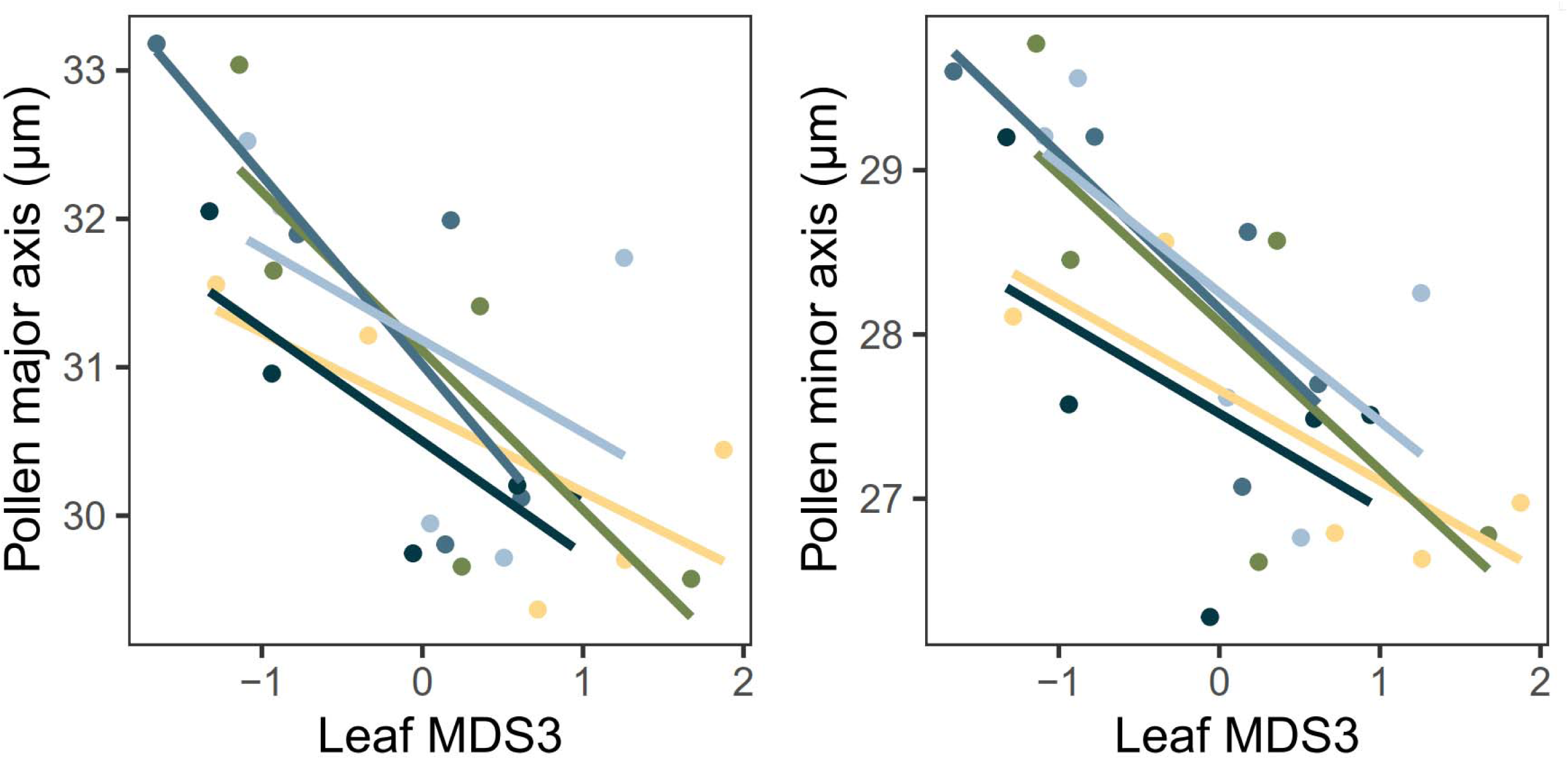
Relationship across branches of individual *Pistacia terebinthus* trees between reduced epigenetic fingerprint of fresh leaves (Leaf MDS3, sample coordinates on the third axis of multidimensional scaling of MS-AFLP dissimilarity matrix, centered to mean zero and standard deviation unit) and major and minor axis length of the pollen grains produced by adjacent inflorescences (see Table 3 for statistical analysis). Each symbol corresponds to a different tree x branch combination, lines are within-tree linear regressions, and color codes denote the five trees sampled.

**Table 3.**
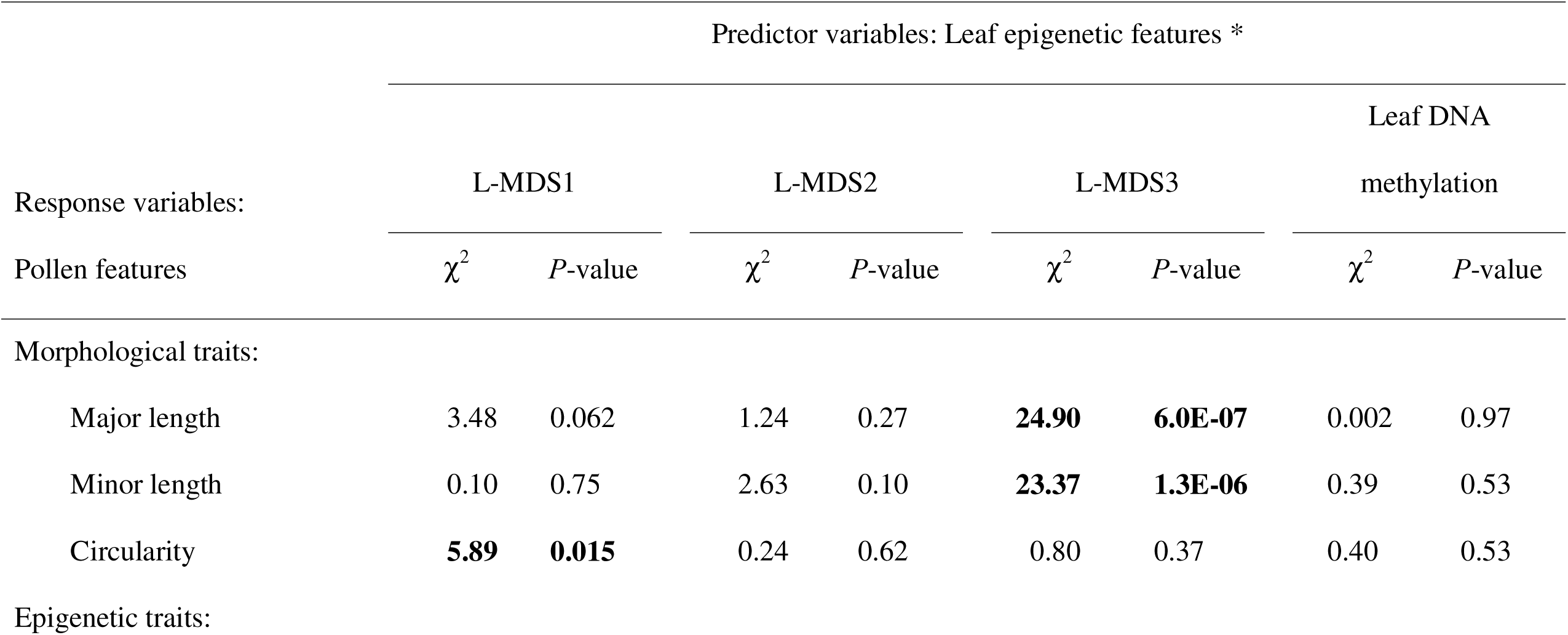
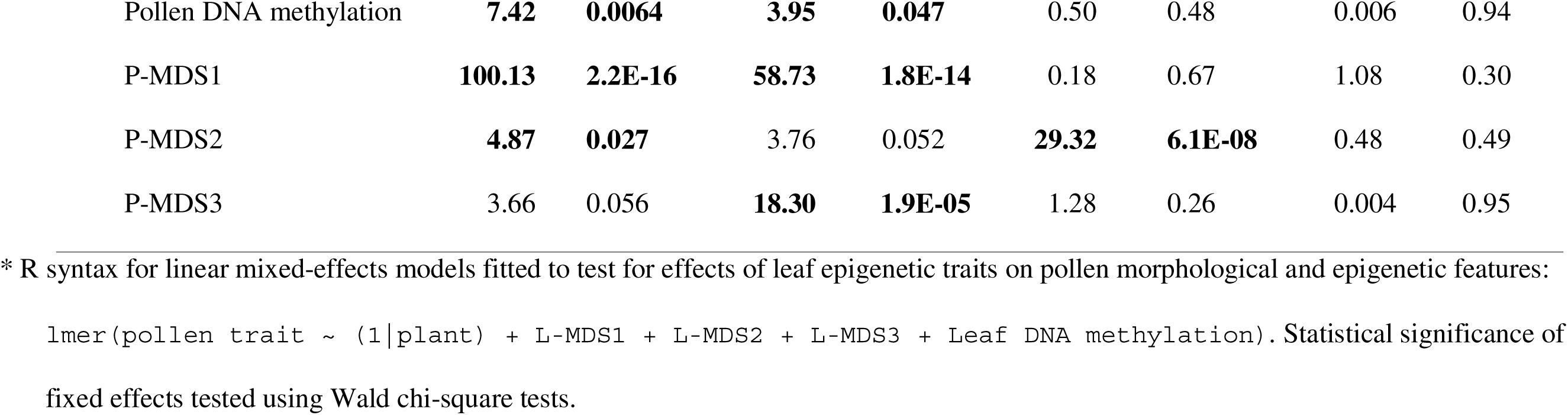
Summary of results of linear mixed-effects models testing for within-tree relationships between morphological 594 and epigenetic variation of the pollen and epigenetic features of adjacent fresh leaves in the same growth axis. L-MDS’s and P-MDS’s stand for multivariate epigenetic fingerprints of leaf and pollen samples, respectively, and are sample coordinates on the three axes obtained by running multidimensional scaling on matrices of pairwise epigenetic dissimilarity between plant x branch combinations with regard to methylation state of MS-AFLP markers. All models included tree identity as a random effect to statistically account for among-tree heterogeneity, but variance components due to this term were small in all cases and have been omitted from the table for simplicity.

Intratree variation in leaf DNA methylation level was not significantly related to any of the descriptors of within-tree epigenetic variation of pollen (Table 3). In sharp contrast, within-plant, among-branch variation in reduced axes scores of multivariate epigenetic features of pollen was largely predicted by the within-plant variation in reduced axes of epigenetic features of leaves, as shown by the five (out of the 9 possible ones) statistically significant associations between the epigenetic axes of leaf (L-MDS1, L-MDS2, L-MDS3) and pollen tissue (P-MDS1, P-MDS2, P-MDS3) across branches of the same tree (Table 3). Two examples of these relationships are illustrated in Figure 3. Branch scores on the first and second reduced axis of epigenetic pollen variation (P-MDS1 and P-MDS2, respectively) were positively related across branches of the same tree to the corresponding scores on the first and third reduced axes of epigenetic leaf variation (L-MDS1 and L-MDS3, respectively). The relationships were closely similar across trees.

**Figure 3.**
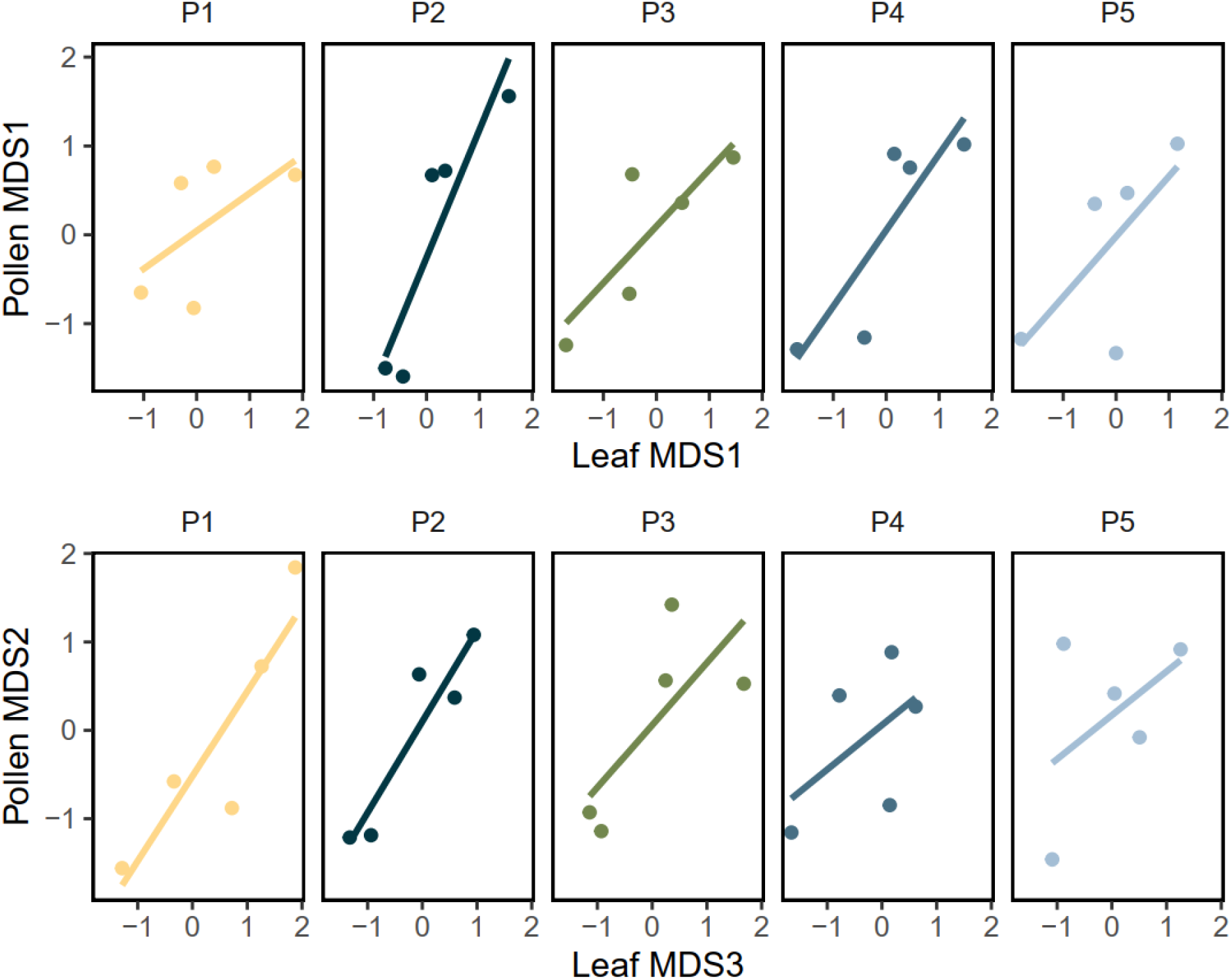
Relationships across branches of individual *Pistacia terebinthus* trees (P1 to P5) between reduced epigenetic fingerprints of fresh leaves and the pollen grains produced by adjacent inflorescences. Pollen MDS1 and MDS2, and Leaf MDS1 and MDS3, are branch coordinates on axes of nonmetric multidimensional scaling of the MS-AFLP dissimilarity matrix for pollen and leaf tissue, respectively (centered to mean zero and standard deviation unit). See Table 3 for results of statistical analyses.

## DISCUSSION

This study has found that *P. terebinthus* trees were internally heterogeneous with regard to both genome-wide DNA cytosine methylation and multivariate MS-AFLP fingerprints of leaf tissue. Estimated variance in epigenetic features of leaves borne by different branches of the same tree exceeded estimated among-tree variance, which was quantitatively minor. In addition, morphological and epigenetic features of pollen grains differed significantly among branches of individual trees. Examples of somatic epigenetic mosaicism in which different homologous parts of the same genetic individual differ in patterns or extent of DNA methylation have been previously reported for clonal (Gao et al., 2010; Bian et al., 2013; Spens & Douhovnikoff, 2016) and non-clonal plants (Das & Messing 1994; Bitonti et al., 1996; Alonso et al., 2018; Herrera et al., 2021, 2022). We are not aware, however, of any previous study documenting epigenetic divergence among the populations of haploid gametophytes produced by different parts of the same plant, as found here for male *Pistacia terebinthus* trees (but see, e.g., Colombo et al., 1983; Wronska-Pilarek & Jagodzinski, 2012; Strelin et al., 2025, for examples of broad subindividual variation of pollen phenotypes in long-lived species; and Maletskii et al., 2008 for evidence suggesting epigenetically-driven subindividual variation in pollen viability). Taken together, our results support the view that somatic epigenetic mosaicism within adult plants of long-lived species is concordant with epigenetic variation in the subsequent gametophytic generation, as implicitly postulated by the epigenetic mosaicism hypothesis of intraplant variation (Herrera et al., 2021, 2022). The statistically significant relationships linking somatic epigenetic with gametophytic epigenetic and phenotypic variation across branches of the same tree (Table 3, Figures 2 and 3) can be parsimoniously interpreted as a cause-effect link between these two layers of intraplant variation implying resemblance by descent. This interpretation is supported by the examples reported in recent years of transgenerational inheritance of epigenetic marks from parent plants to sexually produced F1 progeny and beyond (Bošković & Rando 2018; Rogo et al. 2026; Steadman et al. 2026), all of which must evidently have involved a transitional sporophyte-to-gametophyte transmission. The considerable, albeit imperfect coupling of within-tree variation in epigenetic fingerprints of leaves and pollen (Figure 3) actually suggests substantial inheritance of epigenetic marks from somatic to gametophytic tissues in the same branch, but our data do not allow for a formal test. Results do suggest, however, that whatever the extent of epigenetic resetting taking place during male gametogenesis in *P. terebinthus* trees, it was clearly insufficient to epigenetically homogeneize the gametophyte populations produced by epigenetically distinct parts in the same individual tree. In addition, the resulting phenotypic mosaicism in size and shape of pollen grains represents a possible selective scenario whereby somatic epigenetic heterogeneity within trees can eventually translate into differential male fitness of different branches of the same individual mediated by size-related, epigenetic variation of pollen grains.

In adults of long-lived woody plants such as *P. terebinthus*, within-plant epigenetic heterogeneity of somatic tissues most likely reflects the combination of a fast-ticking epigenetic molecular clock generating stochastic variability which eventually propagates over the branching topology of individuals (Herrera et al., 2021; Yao et al., 2021, 2023; Gardner et al., 2023; Vo et al., 2024), and localized epigenetic changes induced by environmental factors which can act differentially on different sectors of individual plants (e.g., herbivory, light regime, temperature, insolation; Roslin et al., 2006; Herrera & Bazaga, 2013; Zonner & Renner, 2015; Eisenring et al., 2021; Emmerson et al., 2025). Regardless of its proximate origin, intraplant epigenetic heterogeneity can have short-term ecological consequences for the genetic individual, including enhanced within-plant variance in inflorescence and seed size, floral features, progeny performance and impact of herbivores (Alonso et al., 2018; Herrera et al., 2019, 2022; Trevizan et al. 2026). In the same vein, the broad within-plant variation in size and shape of pollen grains found here (see also Strelin et al., 2025) will most likely promote variability in the pollination success achieved by individual *P. terebinthus* trees, since intraspecific variation in pollen grain size can be correlated with germination rate, pollen tube growth and siring success (Cruzan, 1990; Kelly et al., 2002; McCallum & Chang 2016). Epigenetic reprogramming occurring during fertilization (Gutierrez-Marcos & Dickinson, 2012; Kawashima & Berger, 2014; Jo & Nodine, 2024) could shrink the intraindividual variability in pollen epigenetic features found here. It seems unlikely, however, that such additional epigenetic resetting would obliterate the extensive epigenetic variation among pollen of different branches in a tree which had “survived” a thorough reprogramming during gametogenesis.

Recent discovery of fast-ticking evolutionary epigenetic clocks in plants has opened unexplored research avenues at the interface between ecology and evolutionary biology (Yao et al., 2023), particularly in relation to the fate and implications of random epigenetic variants created by such clocks within the confines of otherwise genetically homogeneous individuals (Herrera et al., 2021, 2022). Somatic epigenetic heterogeneity acquired over a parent plant’s lifetime via random epimutations could translate into epigenetic and/or phenotypic differences in its sporophytic descendants (Herrera et al., 2021; Yao et al., 2021, 2023; Shah, 2022; Gardner et al., 2023; Chan et al., 2024; Vo et al., 2024). Our results for *P. terebinthus* provide compelling evidence that intraindividual epigenetic heterogeneity in vegetative parts cascaded into epigenetically heterogeneous haploid gametes. This result confirms in a within-plant context the sporophyte-to-gametophyte transmission of epigenetic marks which takes place at the whole plant level (Takeda & Paszkowski 2006; Herrera et al., 2013), illustrate one proximate mechanism whereby subindividual epigenetic variants can become established in subsequent generations of diploid sporophytes, and suggests some novel hypotheses for epigenetic research on wild populations of non-model plants: (1) Intraplant variation in epigenetic features of pollen grains suggests functional non-equivalence of pollen produced by different branches in the same tree, a possible confounding factor which should be taken into consideration in studies of variation in pollen performance or progeny fitness besides individual genetic background or somatic genetic mutations (Cruzan et al., 2022); (2) the sporophyte-to-sporophyte transgenerational fecundity effects of parental epigenetic heterogeneity previously reported for other plants (Herrera et al., 2019, 2022) could actually be mediated by heterogeneity in phenotypic and epigenetic quality of male gametes; (3) in a random epigenetic clock scenario, intraplant variance in pollen epigenetic traits will tend to increase with age, leading to age-related intraplant variability in pollen phenotype, viability and performance (Strelin et al., 2025); and (4) the inability of epigenetic reprogramming during gametogenesis to completely wipe out somatic epigenetic variability within plants should generate an opportunity for selection on genes which limit or prevent the transgenerational transmission of environmentally-induced epigenetic states (Iwasaki & Paszkowski, 2014). This ultimately implies that some epigenetic variants arising in somatic tissues over individual lifespans could eventually contribute to the phenotypic diversity represented in the next generation of adult plants (Shahzad et al., 2025). By this mechanism, intraindividual epigenetic variation in somatic tissues, arising as a consequence of either localized environmental induction or epigenetic clocks, could eventually contribute to enhance epigenetic and phenotypic diversity in natural plant populations.

## Author contributions

CMH conceptualized the work, conducted field sampling, performed statistical analyses and led the writing; MM developed the pollen purification protocol, prepared pollen samples and measured pollen grains; PB carried out MS-AFLP analyses; CA supervised HPLC analyses, curated results and contributed to interpretations. All authors refined the manuscript and approved the final version.

## Acknowledgments

We thank Mónica Gutiérrez-Rivillo for laboratory assistance, and Consejería de Medio Ambiente, Junta de Andalucía for permission to work in the Sierra de Cazorla and providing invaluable facilities there. Partial financial support was provided by the Ministerio de Ciencia e Innovación, Spanish Government (grant DISTEPIC-PID2022-141530NB-C22).

## Data availability statement

Raw data used in this study are available to editors and reviewers at figshare https://figshare.com/s/940d6a5bcbd924b71c96

